# Galvanotaxis of ciliates: Spatiotemporal dynamics of *Coleps hirtus* under electric fields

**DOI:** 10.1101/2021.12.16.473004

**Authors:** Anna Daul, Marie-Louise Lemloh, Marcel Hörning

## Abstract

Galvanotaxis describes the functional response of organisms to electric fields. In ciliates, the electric field influences the electrophysiology and thus the cilia beat dynamics. This leads to a change of the swimming direction towards the cathode. The dynamical response to electric fields of *Coleps hirtus* has not been studied since the observations of Verworn in 1890 (1). While galvanotaxis has been studied in other cilitates, *C. hirtus* exhibit properties not found elsewhere, such as biomineralization-processes of alveolar plates with impact on the intracellular calcium regulation and a bimodal resting membrane potential, which leads unique electrophysiological driven bimodal swimming dynamics. Here, we statistically analyze the galvanotactic dynamics of *C. hirtus* by automated cell tracking routines. We found that the number of cells that show a galvanotactic response, increases with the increase of the applied electric field strength with a mean at about 2.1 V/cm. The spatiotemporal swimming dynamics change and lead to a statistical increase of linear elongated cell trajectories that point toward the cathode. Further, the increase of the electric fields decreases the mean velocity variance for electric fields larger than about 1.3 V/cm, while showing no significant change in the absolute velocity for any applied electric field. Fully functional galvanotactic responses were observed at a minimum extracellular calcium concentration of 20 *µ*M. The results add important insights to the current understanding of cellular dynamics of ciliates and suggest that the currently accepted model lags the inclusion of the swimming dynamics and the complex calcium regulatory system of the cell. The results of this study do not only extend the fundamental understanding of *C. hirtus* dynamics, but also open possibilities for technical applications, such as biosensors or microrobots in the future.

## Introduction

Galvanotaxis is a directed reaction of organisms to an applied electric field. The phenomenon of galvanotaxis has been observed in multicellular organisms such as fish (2), as well as in unicellular organisms such as bacteria (3), amoebae (4), ciliates and flagellates or different types of eukaryotic cell types (5, 6). It has been shown that galvanotaxis also plays a role in embryonic development and wound healing (5–7). In recent decades, galvanotactic behaviour of protozoa has been applied for sample purification (8, 9), was discussed to be applied for automation of toxicity assays (10) and the phenomenon was investigated in context of bioinspired micro-robots. (11–13). The galvanotaxis of ciliates was first described in the 1890s by Verworn (1, 14) and the reaction of ciliates to the application of a constant current was observed. It was shown that most ciliates exhibit a negative galvanotaxis, which means they swim towards the cathode in the electric field (14–16).

Galvanotaxis is not an intrinsic taxis like phototaxis or chemotaxis, but a reaction caused by external intervention on electrophysiology. In the case of ciliates, the electric field causes a change in the cilia beat, which results in the changed swimming behaviour. According to the current model, the activation of voltage-dependent ion channels in the ciliary membrane by the electric field and the resulting physiological processes are responsible for the galvanotactic response (17). If an electric field is applied to the medium externally, an equipotential surface is formed. This encloses the cell, which can be considered a conductor (18, 19), and creates a potential gradient between the two ends of the cell along the anodic and cathodic direction. Thus, the cathodic facing membrane site depolarizes and the opposite side hyperpolarizes (20). Depending on the local membrane potential, the distinct voltage-dependent ion channels in the ciliary membrane will be opened. At the anodic end, the voltage-dependent potassium channels open up, which leads to an outflow of K^+^ from the cilia resulting in an increase in the beat frequency (21–23). The increase of the beat frequency depends on the degree of polarization (24). At the cathodic side, the voltage-dependent calcium channels are opened, which leads to an influx of Ca^2+^ into the cilia. If the Ca^2+^ concentration in the cilia is increased, there is a rotation of the beat direction until it is completely reversed (19, 25–30). The reversed and amplified ciliary beat leads to a backward and a forward force which compete. Depending on the position of the cell in the field, the reverse stroke appears on a smaller cell surface area. The asymmetry of the forces of cilia at the membrane results in a torque, which causes the cell to orientate itself towards the cathode and consequently to swim in this direction (31).

The ciliate *Coleps hirtus* (Fig. 1) lives planktonic in freshwater and saltwater habitats and feeds omnivorously on algae, bacteria, ciliates, small invertebrates and dead organic material (32, 33). The cell body of *C. hirtus* is 40-65 µm long and 18-35 µm wide, it is barrel-shaped with an almost circular cross section (32). *C. hirtus* is particularly well characterised by its alveolar plates composed of amorphous calcium carbonate and organic material (34, 35) that are arranged in 6 belts with 12-20 longitudinally rows each (32, 33). The cilia of *C. hirtus* are also arranged in 12 to 20 longitudinal rows, protruding outwards from small openings between the alveolar plates and a significantly extended caudal cilium located at the posterior end (Fig. 1C-D) (32, 33, 36). *C. hirtus* shows an interesting swimming behaviour of alternating periods of straight forward swimming in an counter clockwise hellical spiral (32) and localized tumbling-like circular swimming, which is related to a Ca^2+^ dependent plateau potential (37).

**Fig. 1.**
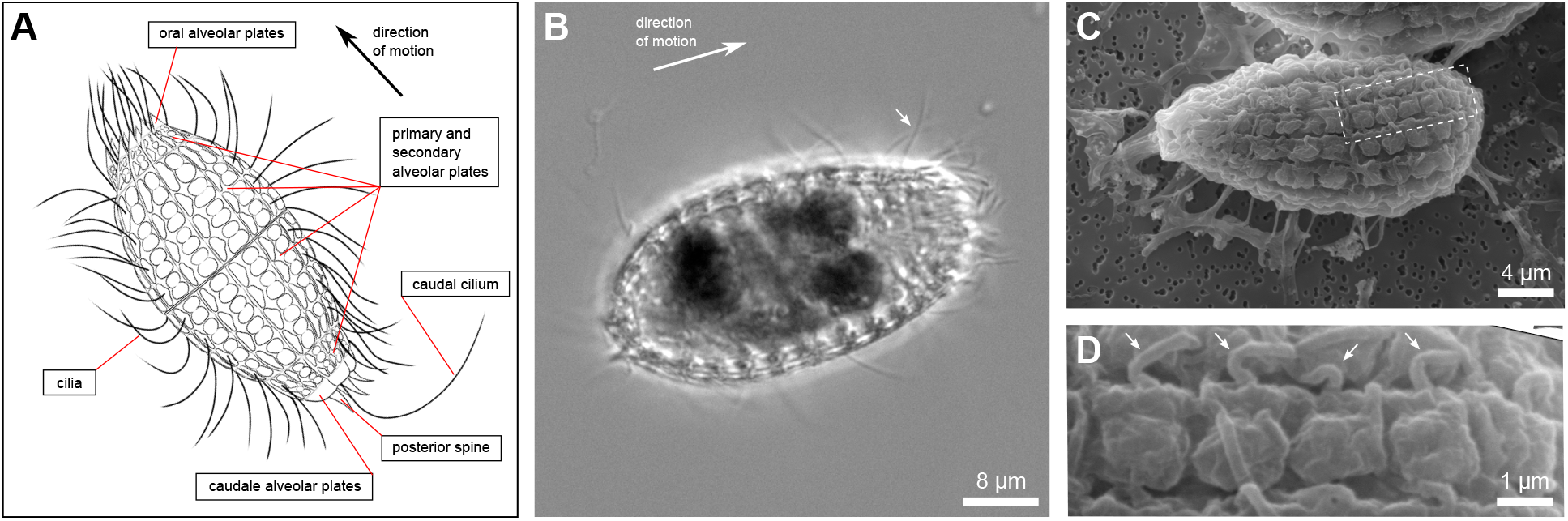
General morphology and whole cell visualization of *C. hirtus*. (A) Schematic drawing, (B) Light microscopic image and (C, D) Scanning electron microscopic image of *C. hirtus*. A high-speed movie of one cell is shown in Supplemental Materials.

The galvanotactic behaviour *C. hirtus* has not been scientifically documented during the last century. In 1890 Verworn (14) observed that *C. hirtus* swim towards the cathode and the swimming velocity decreases under electric fields. It was also observed that under larger electric fields *C. hirtus* sometimes swim backwards towards the anode (14). The aim of this work is to revise and improve those earlier observations by using state of the art observation techniques and image analysis methods. In particular the focus is on identifying the galvanotactic response and spatiotemporal swimming dynamics of *C. hirtus* as a function of the electric field strength.

**Results**

*C. hirtus* cells constantly swim to keep feeding by self-propulsion through periodic and synchronized cilian motion. The cilia are arranged on the cell surface covered by the plasma membrane (Fig. 1) and can respond to external electric fields. While the cells do not possess any inherent or intrinsic galvanotaxis, the application of an external electric field leads to changes in the beating dynamics of cilia, which in turn leads to an effective change of the direction of motion toward the cathode. In order to systematically observe and quantify the dynamics of *C. hirtus* under various external electric fields, a customized 3D-printed chamber was manufactured (Fig. 2A) to setup an observation chamber that enables tracking of the motion dynamics of cells under electric fields up to *E* ≃11V/cm. A common framework is introduced to define the direction of motion *φ*, as illustrated in Fig. 2B. The directions *φ* parallel to the cathode and anode are defined as 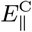 and 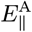, and the directions perpendicular, right- and leftward respectively the cathode are defined as 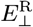 and 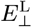. Using this framework we are able to define the direction of motion of single *C. hirtus* cells (see Materials and Methods), as exemplarily shown for three cells at different applied electric field strengths (Fig. 2C). Individual cells are observed for 20 seconds and their spatiotemporal traces are linear color-coded by the observation time from the start at *t*_0_ = 0 to the end *t*_end_ = 20s. Visual inspection of single traces at different electric fields revealed an apparent stabilization of swimming helicallity and an increase in swimming directionality toward the cathode (Fig. 2C, *E* = 4V/cm), e.g. an decrease of the angle difference Δ*φ*. The latter is defined as the angle difference between 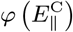 and the cell’s direction of motion *φ*_cell_, which was calculated from the first **r**_0_(*x*_0_, *y*_0_, *t*_0_) and last **r**_end_(*x*_end_, *y*_end_, *t*_end_) observed cell position, as

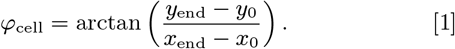

**Fig. 2.**
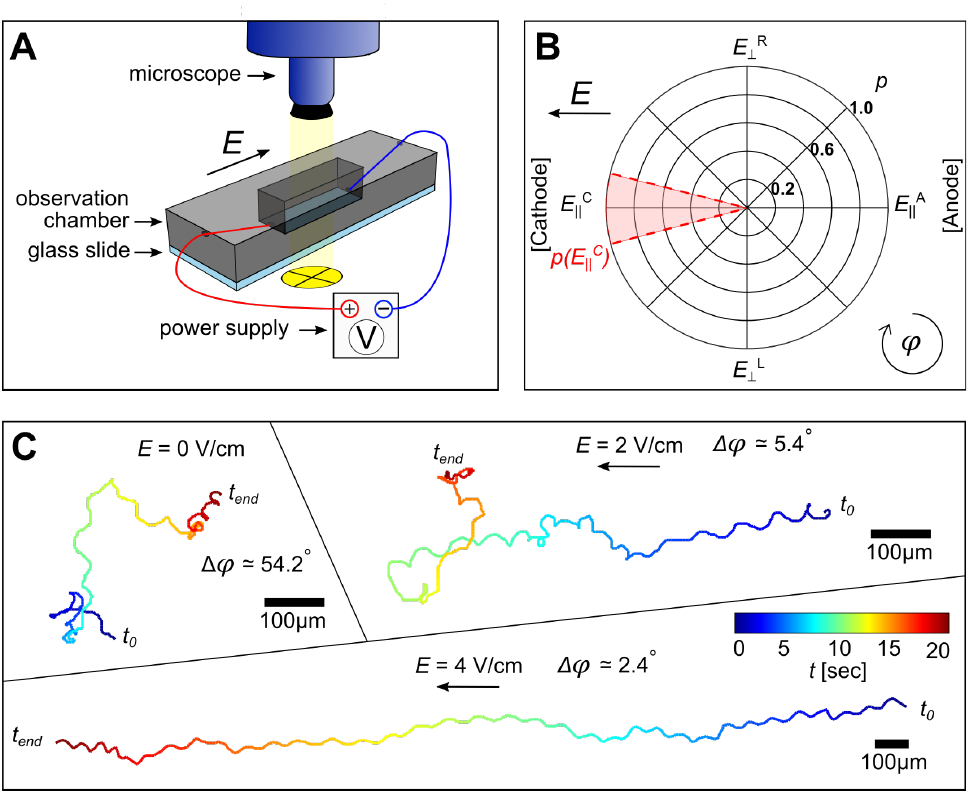
Setup and dynamics quantification strategy of *C. hirtus* under influence of an electric field. (A) Scheme of the experimental setup using a custom-designed 3D-printed imaging chamber with cable-inlets to apply electric fields of up to *E* = 11 V/cm (see Materials and Methods). (B) Quantification scheme to analyse the motional direction of *C. hirtus* relative to the electrodes. The direction is categorized by the movement of cells parallel to the electric field toward the cathode 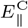 and anode 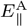, as well as the left and right handed direction perpendicular to the cathode as 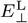 and 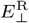, respectively. The probability 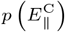 for cells moving toward 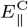 with an arc span of 0.2*π* is exemplarily highlighted in red color. (C) Three *C. hirtus* cell-traces exemplarily observed at different electric fields *E* = 0 V/cm, *E* = 2 V/cm and *E* = 4 V/cm. The traces are color-coded by the observation time *t*. The start and end of each trace is marked by *t*_0_ = 0 and *t*_end_ = 20 s. The black arrows indicate the direction of the applied electric field.

### Impact on cells under repeated electric fields

For statistical quantification of the motional characteristics under electric fields the cell number was increased. This enabled the observation and tracking of *C. hirtus* cells simultaneously under comparable conditions. Figures 3A and B show traces of cells without (*E* = 0) and with an applied electric field (*E* = 8V/cm). At *E* = 0 cells show helical swimming dynamics erratically interrupted by sudden changes in helicallity and changes of direction (see also Fig. 2C). Some ciliates barely swim and rotate on the spot. With an applied electric field almost all cells swim toward the cathode 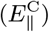 (Fig. 3B).

**Fig. 3.**
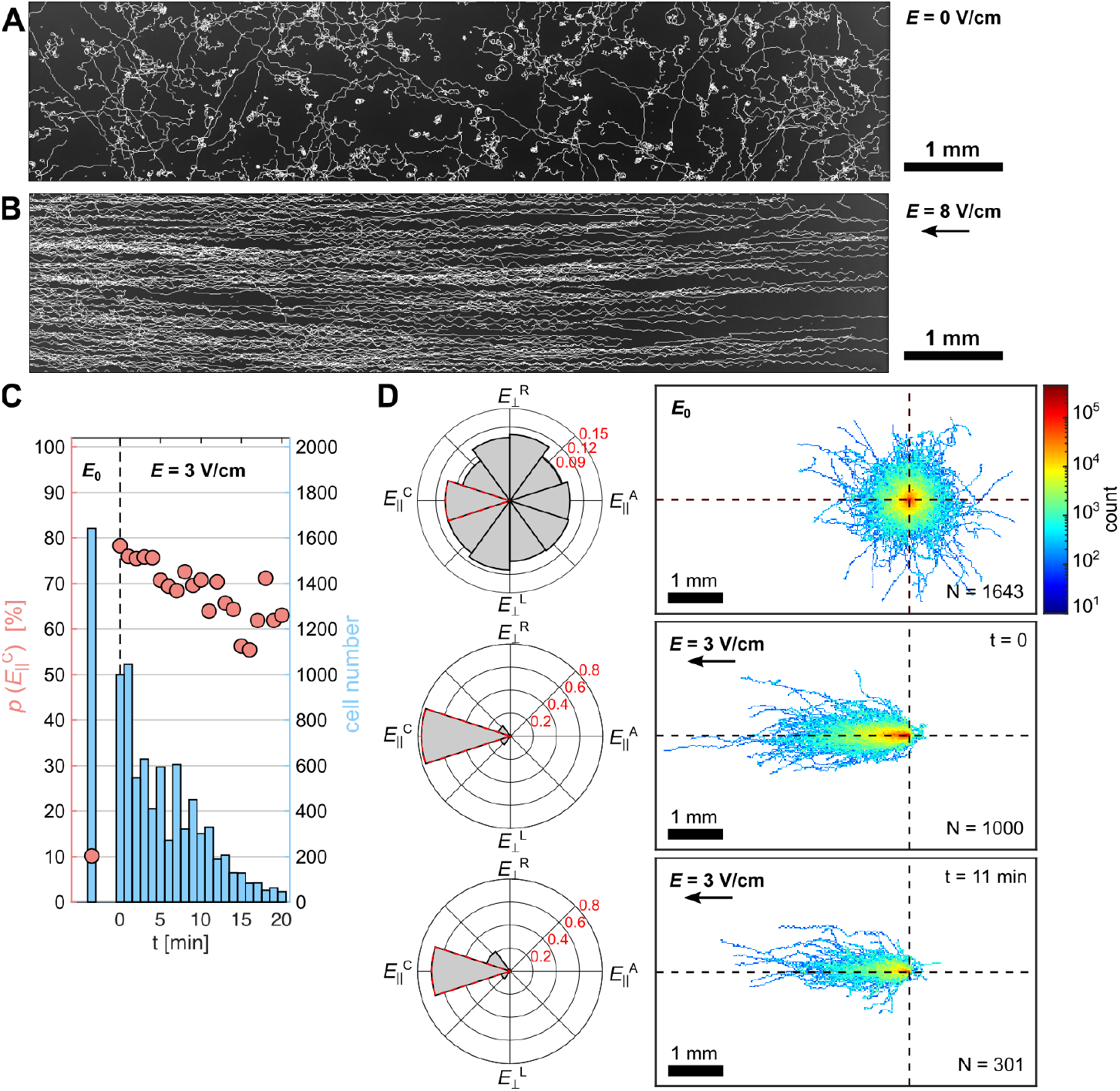
Quantification of the direction of motion of *C. hirtus*. (A) and (B) show overlaid trajectories for cells without (*E* = 0 V/cm) and with (*E* = 8 V/cm) the influence of an externally applied electric field. (C) Statistical evolution of the probability 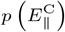 that indicates the fraction of cells moving toward the cathode (red circles). The dashed line at *t* = 0 indicates the start of the electric field application (*E* = 3 V/cm). Cells were observed every minute for 20 seconds by periodic switching the direction of the applied electric field. Blue bars indicate the respective number of observed cells. (D) Statistical analysis of cells without (*E* = 0 V/cm) and with (*E* = 3 V/cm) an electric field at *t* = 0 and *t* = 11 min. Left are shown the radial probability density distributions and right the overlaid cell traces centered at the same initial spatiotemporal position (*t*_0_, *x*_0_, *y*_0_), as indicated by the dashed lines. The variable N indicates the number of analyzed cells.

To monitor the effect of repeated experiments under electric field applications, we observed cells first without electric field (*E* = 0) followed by subsequent repeated observations of the same cell population at *E* = 3 V/cm for 20 s. The electric field direction (position of anode and cathode) was switched periodically every minute to ensure a sufficient amount of cells remained within the field of view. The observation was continued 20s after the electric field was switched giving the cells sufficient time to respond to the change. The probability of cells moving toward the cathode 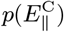, defined in the range of 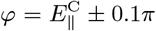 (Fig. 2B) was mapped over a duration of 20 min (Fig. 3C). Without an electric field about 10% of cells moved toward 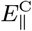, as theoretical predicted for a non-directional isotropically moving cell population. Once the electric field is applied (*t* = 0) most of the cells 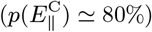 moved toward the cathode 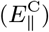. After about 6 min 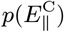 (6 observations) the amount began to decrease constantly down to about 60% after 20 min. Further, the number of cells tracked within the observation window dropped exponentially (blue bars, Fig. 3C). Figures 3D shows three probability density graphs that quantify the direction of motion for *E* = 0 and *E* = 3 V/cm at *t* = 0 and *t* = 11 min. The two-dimensional cell trace histograms are shown on the right by defining the initial cell position at the intersection of the two dashed lines. Although, an observation time of up to 5 minutes would be reasonable without inducing significantly measurable cell exhaustion or damage, further investigations were performed using untreated cells with an observation time of 20 seconds only. This ensures the reproducibility of the statistical outcome.

### Quantification of motion dynamics

For the quantification of the direction of motion *φ* all *C. hirtus* cells were tracked and analysed. The probability distribution of the direction of motion *p*(*φ*) was quantified depending on the electric field strengths *E* (Fig. 4A). The latter were varied between *E* = 0 V/cm and *E* = 11 V/cm in steps of Δ*E* = 0.2 V/cm up to *E* = 3 V/cm followed by steps of Δ*E* = 1.0 V/cm. The direction of motion was calculated for all cells and plotted as normalized histogram for each probed *E*. Figure 4B shows the probabilities 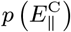 (blue circles) and 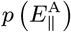 (red squares) as a function of *E*. A monotonic increase from 10% up to about 90% was observed for 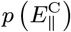, while 10% indicates the non-directional isotropically distributed cell motion when no electric field is applied (see also Fig. 3D, *E* = 0 V/cm). The data are fitted to normal cumulative distribution functions (Eq. 7) indicating that the galvanotactic response of *C. hirtus* cells is normal distributed at *Ē*_*p*_ = 2.1 ±1.0 V/cm. A small fraction of cells (1%-2%) exhibited a different swimming direction toward the anode as indicated by 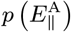 for *E* ≳3V/cm and visually apparent in Fig. 4A at 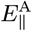. At the highest applied electric field *E* = 11 V/cm about one out of 20 cells move toward the anodal direction when considering the two possible directions to the anode 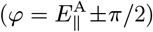 and cathode 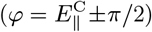 only.

**Fig. 4.**
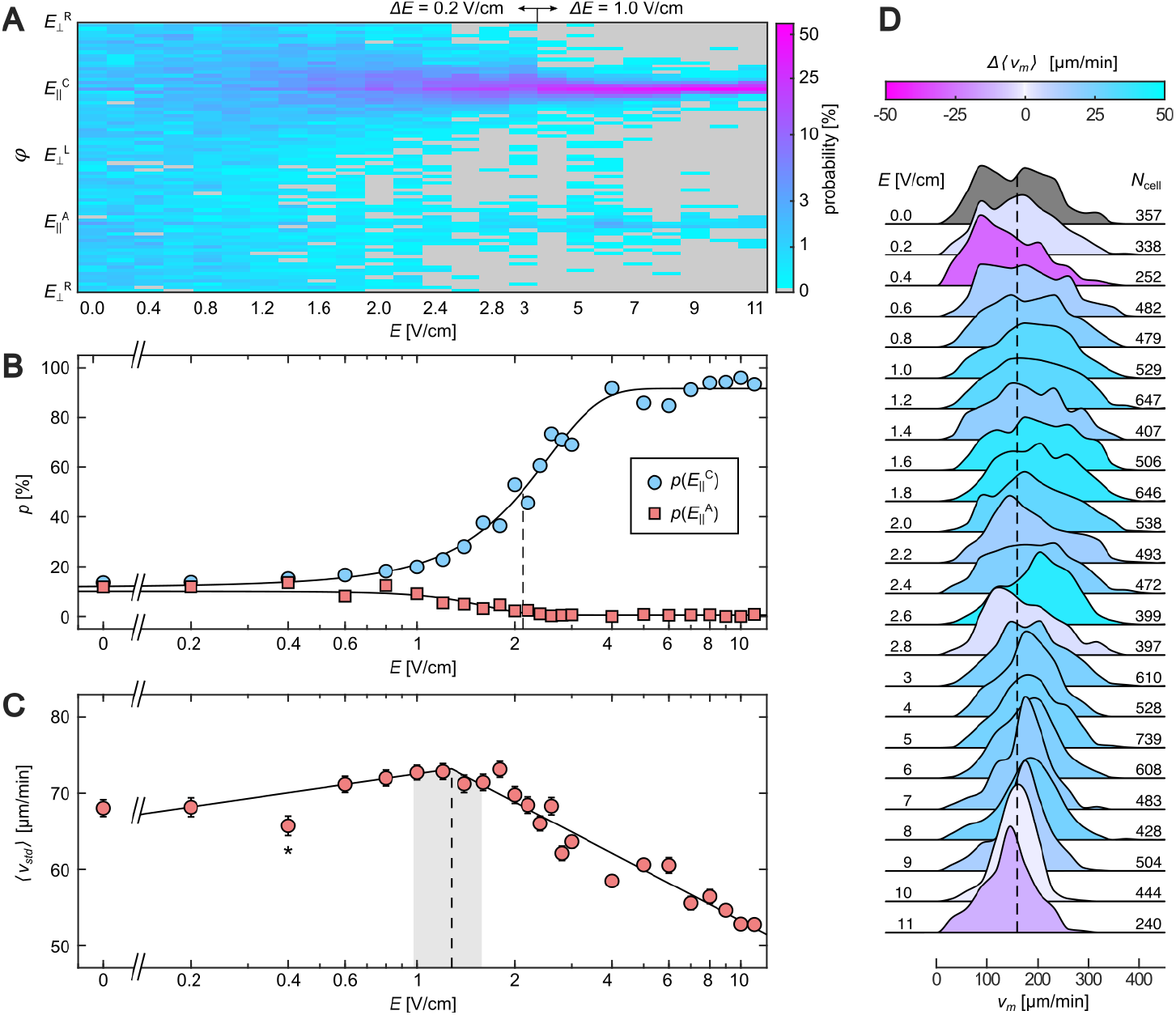
Quantification of spatiotemporal dynamics of *C. hirtus*. (A) Probability density distribution of the direction of motion under different applied electric fields. (B) Probabilities 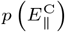 (blue circles) and 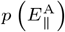 (red squares) as a function of *E*. The black solid lines are the respective least-square fits using a normal cumulative distribution functions (see Eq. 7) with indicated mean (dashed line) and standard deviation as *Ē*_*p*_ = 2.1 *±* 1.0 V/cm. (C) Mean velocity variability ⟨*υ*_std_⟩ as a function of *E*. Errorbars show the standard error. The black solid line is the least-square fit of piecewise logarithmic functions (see Eq. 8) with indicated breakpoint (dashed line) and 95% confidence interval (grey range) as *Ē*_*v*_ = 1.3*±* 0.3 V/cm. The asterisks indicate the data points that were omitted for the fit. (D) Normalized velocity distributions of *C. hirtus* for different applied electric fields. The color scheme indicates the respective difference to the median velocity at *E* = 0 V/cm shown in grey color on the top.

Next, we quantified the velocity variability by calculating the mean and standard deviation *v*_m_ ± *v*_std_ of each cell. From that the mean velocity variability ⟨*v*_std_⟩ for each applied electric field was obtained using a log-transformation and the Finney estimator approach (39) (see Material and Methods). Figure 4C shows ⟨*v*_std_⟩ as a function of *E*. The data indicate a slight increase of ⟨*v*_std_⟩ until *Ē*_*v*_ ≃ 1.3 V/cm, which indicates the breakpoint of the least-square fitted piecewise logarithmic function (Eq. 8). Larger electric fields lead to a significant decrease of ⟨*v*_std_⟩ that corresponds to the stabilization of swimming helicality, as observed at the single cell traces (Figs. 2C and 3A-B). While ⟨*v*_std_⟩ significantly depends on *E*, we did not observe a correlation to the mean velocities of cells (Fig. 4D, Fig. S1). Hence we conclude that the electric field reduces the velocity variability that leads to an increase in monotonic swimming directionality without changing the mean velocity of cells.

An independent verification of this result is the correlation between ⟨*v*_std_⟩ and cell trajectory. The latter was quantified by fitting an ellipse to the trajectory of each individual cell, which can take on values between 0 (strongly elliptic) and 1 (circular), see inset of Figure 5. From the aspect ratio of the ellipse *r*_E_ = *minoraxis* / *majoraxis* the mean elliptic aspect ratio ⟨*r*_E_⟩ was caltulated from all observed cells. Figure 5A shows ⟨*r*_E_⟩ as a function of *E* with a fit of a normal cumulative distribution function (see Eq. 7). Similar to the probability 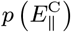 (see Fig. 4B) is the galvanotactic response of *C. hirtus* cells normal distributed at *Ē*_*r*_ = 2.6 ±1.1 V/cm. The comparison between ⟨*r*_E_⟩ and ⟨*v*_std_⟩ led to a linear relation with the mean slope of 0.02 min/*µ*m (Fig. 5B). Contrarily, the mean velocity ⟨*v*_m_⟩ did not show any significant dependence to ⟨*r*_E_⟩ (Fig. 5C). The linear relation between ⟨*r*_E_⟩ and ⟨*v*_std_⟩ is also visually inspectable by the distributions of the elliptic aspect ratio and velocity variability (Fig. 5D). Low electric fields (*E* ≤ 1 V/cm) lead to equally distributed *r*_E_ and larger *v*_m_ that changes gradually with an increase in *E*. At the largest applied electric field *E* = 11 V/cm most of cells have a low *r*_E_ close to zero and a lower velocity variability ⟨*v*_std_⟩ ≃ 52*µ*m/min (log ⟨*v*_std_⟩ ≃ 3.9*µ*m/min), indicating long directed cell trajectories with constant helicity and therefore less variation in velocity.

**Fig. 5.**
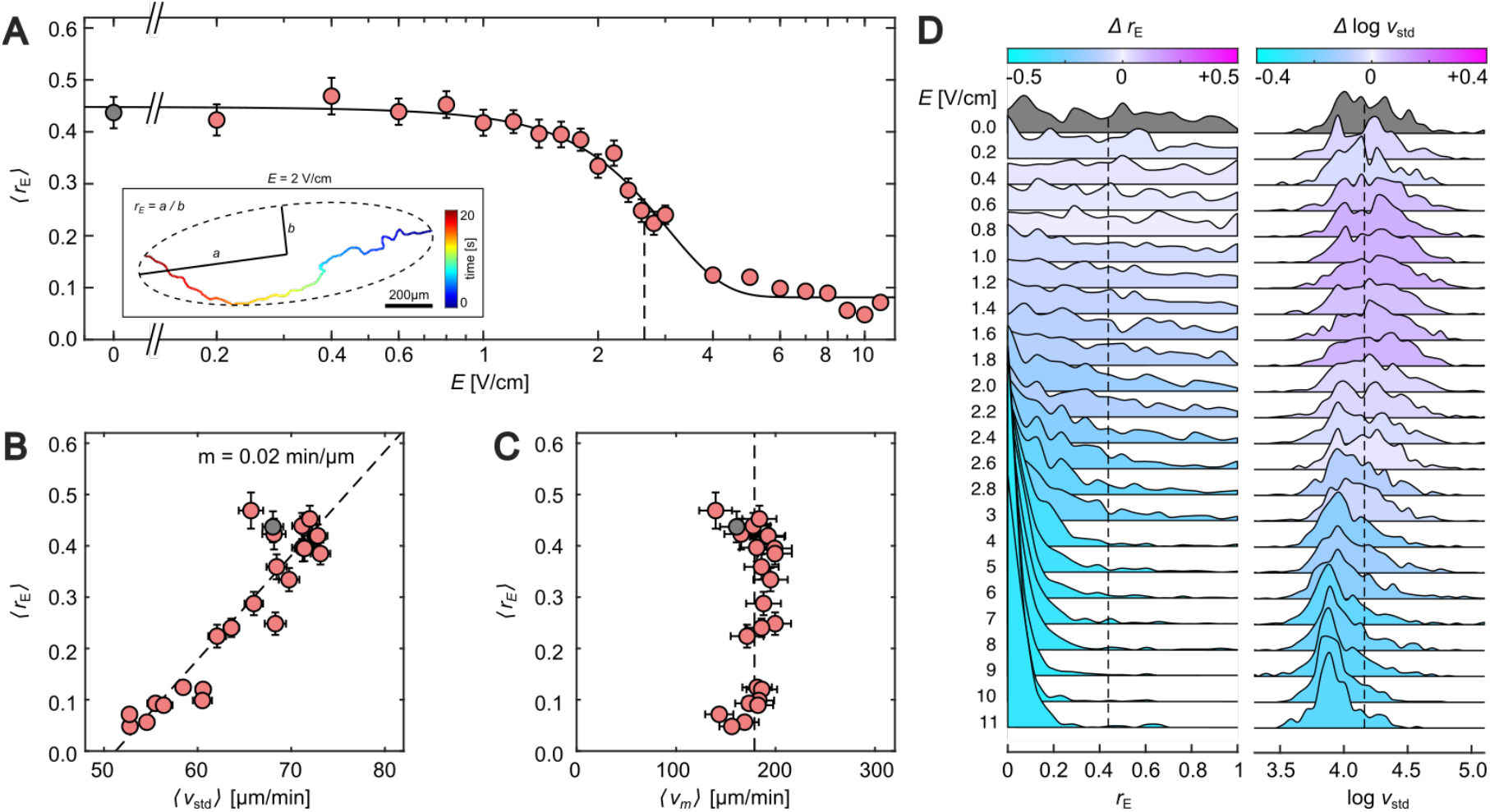
Correlation analysis of spatiotemporal cell-trajectory dynamics. (A) Mean elliptic aspect ratio ⟨*r*_E_⟩as a function of *E*. The black solid line is the least-square fit of a normal cumulative distribution function (see Eq. 7) with indicated mean (dashed line) and standard deviation as *Ē*_*r*_ = 2 6 *±* 1 1 V/cm. (B) Relation between mean elliptic aspect ratio and mean velocity variability. The dashed line indicates the linear fit with the slope *m* = 0.02 min/*µ*m. (C) Relation between mean elliptic aspect ratio and mean velocity. The dashed line indicates the average of the mean velocity. (D) Normalized elliptic aspect ratio (left) and velocity variability (right) distributions of *C. hirtus* trajectories for different applied electric fields. The color scheme indicates the respective difference to the medians at *E* = 0 V/cm shown in grey color on the top of each distribution. Errorbars show the standard error.

### Calcium concentration dependence

Since calcium is essential to regulate mobility and swimming behaviour of *C. hirtus* cells, the role of the extracellular calcium concentration *C*_Ca_ in the culture media was investigated with respect to the galvanotactic behavior of the cells. The cells were observed for 20 seconds without or with a moderate electric field of 3 V/cm that is close to the calculated mean electric fields, *Ē*_*p*_ and *Ē*_*r*_. The over all direction of motion for both recordings were quantified following the same approach as shown in Figs. 4A and B. Figure 6A shows the normalized histograms depending on *C*_Ca_ with and without applied electric field. For all calcium concentrations a preferred direction of motion toward 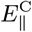 is observed with a visibly narrower distribution for Calcium concentration of 20 *µ*M or more, while no preferred direction of motion is observed without an electric field. Figure 6B shows the calculated probabilities 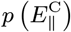 as a function of Ca^2+^ or cells with (*E* = 3V/cm) and without (*E* = 0V/cm) applied electric field (see details at Materials and Methods). Without calcium in the medium, the majority of ciliates do not move. The few that do move show a reduced responds 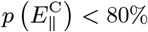 to the applied electric field. A minimal calcium concentration of *C*_Ca_ = 20*µ*M was necessary to observe a normal responds 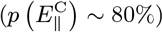 to the electric field similar as observed in standard medium (*C*_Ca_ ≃ 63*µ*M, dashed line). For the mean velocity variation ⟨*v*_std_⟩ only a minor difference between cells with and without electric field was observed for Ca^2+^ *>*≃ 60*µM*, comparable to the results shown in Fig. 4C. A more significant change was observed for the changes in cell trajectories (Fig. 6D). Again only at a minimum *C*_Ca_ of 20 *µ*M an increase of length in cell trajectory (decrease in ⟨*r*_E_⟩) from about ⟨*r*_E_⟩ ≃ 0.45 at *E* = 0 to about ⟨*r*_E_⟩ ≃ 0.2 at *E* = 3 V/cm was observed. These results confirm our previous findings that an externally applied electric field increases the directionally of the cell motion dynamics by reducing the velocity variability leading to a more constant and effective cell motion toward the cathode.

**Fig. 6.**
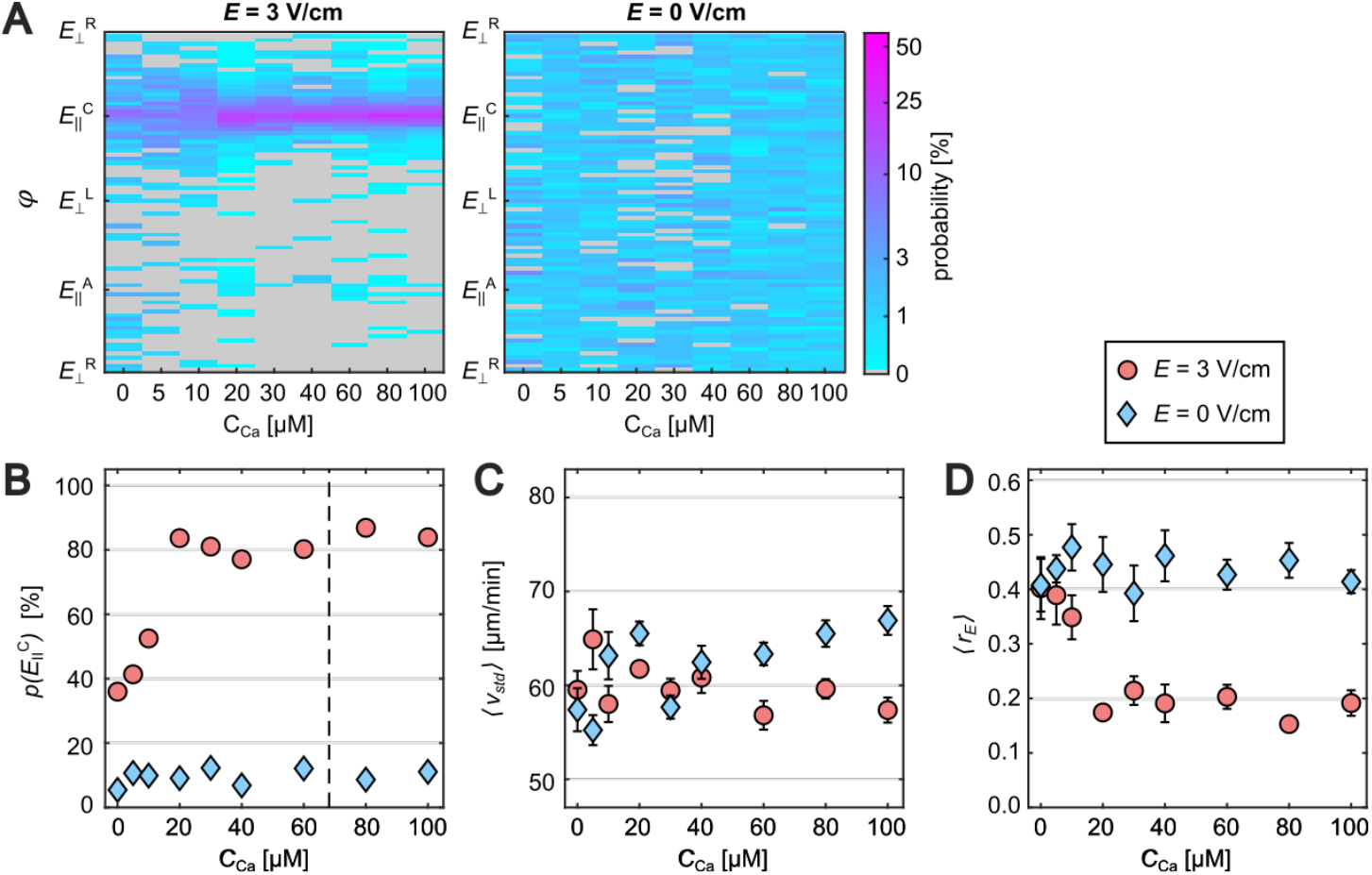
Influence of extracellular calcium. (A) Probability density distribution of the direction of motion at different extracellular concentrations of calcium *C*_Ca_ with (*E* = 3 V/cm) and without applied electric field. (B) Probability 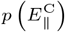as a function of *C*_*ca*_. (C) Mean velocity variability ⟨*v*_std_⟩ as a function of *C*_ca_. (D) Mean elliptic aspect ratio ⟨*r*⟩ as a function of *C*_Ca_. Red circles and blue diamonds indicate the values with (*E* = 3 V/cm) and without applied electric field. Errorbars show the standard error. The dashed line indicates the *C*_Ca_ ≂ 68*µ*M as used in this study.

## Discussions

While galvanotaxis of ciliates has been studied in recent decades, with particular focus on *Paramecium* sp., *Coleps hirtus* have not been studied in a quantitative manner (14). In this work, for the first time the gavanotaxis of *C. hirtus* was statistically and quantitatively analysed using state of the art methods. The characteristic motion dynamics were quantified by large population of *C. hirtus* to obtain statistical measures.

In the absence of an electric field, the expected swimming behaviour could be observed in a customized observation chamber. As soon as an electric field was applied, the direction of motion of ciliates aligned along the field towards the cathode. The proportion of cells swimming parallel to the electric field increased with increasing electric field until saturation was reached. A similar cell response has been observed in the ciliate *Tetrahymena vorax* before (14). The cellular behaviour is thereby explained in such a way that at field strengths below the saturation point, the reversed ciliary beat on the cathodal side is weaker than the amplified cilia beat on the anodal side of the cell. As a result, the torque is not sufficient for all cells to ensure complete alignment in the direction of the cathode. Interestingly, we could show twice indepentendly, by analysing the velocity variations and trajectory quantification that the majority of the cells swim towards the cathode at a value of about 2 V/cm. It remains unclear whether an individual cell needs to experience a defined cell-specific minimal electric field strength to to respond to the electriic field. However, more likely is that the increase in electric field strength leads to an increase in cell response. Although speculative, this could be caused by the constantly changing membrane curvature that is facing the electric field. The electric field threshold might be a function of the local membrane curvature in *C. hirtus*, similarly as shown in conducive cardiac tissue with small intercellular clefts (40). There the degree of curvature changes the critical electric field to induce an increase in ionic-current to excite an action potential (41). Thus larger electric fields may stabilize the induced ionic inflow, because the applied electric field can induce intracellular ionic currents at a wider range of membrane curvatures that face the electric field, and therefore keeping the direction of motion more constant. This hypothesis is likely true considering the similarity of the physical parameters of *C. hirtus* and cardiomyocytes, such as membrane resting potential (∼ −50 mV), membrane resistance (∼ 1*G*Ω), electric field strength (∼ 1 V/cm) and membrane curvature (∼ 1*µm*^−1^), leaving the specific membrane organisation aside. Other factors such as phase of the cell in the cell cycle may also have an effect on the sensitivity of the cell to electric fields. As observed in the past, a small proportion of cells < 5% swim backwards to the anode at the strongest electric fields (14) but have now been quantified statistically for the first time. This phenomenon has also been observed in other ciliates such as *Paramecium* sp. (42) and *Tetrahymena vorax* (43). Since these ciliates were usually described as swimming slowly backwards, the observations were explained in such a way that with stronger electric fields, depolarisation causes the reverse beat to dominate the forward beat which leads to the cell swimming slowly backwards in the direction of the anode.

Unlike other studies, we did not look at individual cells, but at the response of a larger group of cells. This allowed us to identify and quantify important characteristics of the galvanotactic behaviour of *C. hirtus* and to create the basis for further studies, e.g. to refine physical models for galvanotaxis (31). The newly developed experimental set-up enabled the observation of motion response of a large number of cells to examine the distribution of speed and cell trajectorines at a wide range of electric field strength. In contrast to prior observations (14), we did not observe a decrease in cell speed, but a reduction of the variance of speed. This can be attributed to an influence of the galvanic field on the membrane potential. The periodic change between two different resting potentials, which cause the special bimodal swimming pattern of *C. hirtus* (37), no longer occurs. Thus, it can be said that first the electric field leads to a more constant mean velocity of the individual cells and second interferes with the mechanisms that regulate the characteristic bimodal swimming behavior of *C. hirtus* consisting of alternating forward and circular swimming pattern. The results indicate the importance of extracellular calcium for cell dynamics. Without calcium in the medium, a the majority of ciliates do not move. The few that do move show a reduced response to the electric field. A minimum calcium concentration of 20 µM is necessary to observe a normal response to the electric field, similar to the standard medium. However, no influence of higher calcium concentrations could be observed.

For *Paramecium* sp. it is known, that the intracellular calcium concentration can change the direction of the ciliary beat (25, 30), and that blocking different calcium channels and manipulating calcium storages has an influence on the galvanotaxis (44). However in *C. hirtus* studying intracelluar calcium dynamics is challanging, because of the numerous different calcium channels in ciliates, the alveolar vesicles as calcium storage sides and, in case of *C. hirtus* amorphous calcium carbonate containing alveolar plates. The experimental set-up and the advanced image analysis will enable the development of a refined galvanotaxis model in further studies. While these results do not contradict the previous galvanotaxis model (17, 31), they do indicate that the model is not yet complete, as the current model does not include all regulatory elements, e.g. the inclusion of species specific swimming behavior and the complex calcium regulation system of the cell. In context of bioinspired microswimmers, using ciliates under galvanotactic control as micro robots is an exciting field of research that has gained importance in recent decades (11, 12). Technically, there are limitations in power supply and wireless control below a certain size. The bimolecular motors of ciliates are easier to handle. Thus, ciliates can be considered as small micromachines that can be controlled by external stimuli with potential applications, in the medical and biological field, or act as sensors for robotic applications. Therefore, cilian beats and the electrophysiology should be considered in future 3D models.

## Materials and Methods

### Cell Culture

*Coleps hirtus* cultures were collected from Trentsee (Plön) in Germany and cultured for several years under controlled conditions in Wright’s cryptophytes (WC) medium (pH 7.6 - 8.0;) in the dark at 20 °C using Erlenmeyer flasks equipped with oxygen permeable screw caps (see supplemental material). Cultures were fed with *Cryptomonas* sp. every two days and flocculations were regularly removed by using a 40 µm filter. *Cryptomonas* sp. (SAG26.80, SAG culture collection Göttingen, Germany) was grown in Desmediacean medium (MiEB12, SAG, http://epsag.uni-goettingen.de) at 20 °C under photoperiodic lightening (12/12 hours light/dark). Prior feeding *Cryptomonas* sp. was centrifuged (4000×g, 4 min; Rotana 460 R; Hettich Zentrifugen) and resuspended in WC medium. Both, *C. hirtus* and *Cryptomonas* sp. cultures were treated under sterile conditions. The experiments were carried out on the same day of the feeding cycle to obtain reproducible conditions. On the experimental day the culture were fed and filtered (40 µm). A total of 1.2 ml of culture was used for one experimental observation. Culture media with different calcium concentrations between *C*_Ca_ = 0 µM and *C*_Ca_ = 100 µM were compared. Cells were transferred to the Ca-free medium before observation to allow the cells to adapt to it. The desired Ca^2+^ concentration was adjusted with a CaCl2 × 2H2O solution.

### Experimental setup

A custom-designed 3D-printed imaging chamber (76 × 26 × 10 mm [LWH]; observation window: 10 × 30 × 10 mm [LWH]; PLA; Prusa) attached to a microscope slide (76 × 26 × 1 mm [LWH] R. Langenbrinck GmbH) was used for the live imaging of cells (Fig. 2A). The microscope slide was permanently glued and sealed to the chamber using PDMS (1:10; SYLGARD® 184; Dow Deutschland Anlagengesellschaft mbH). For the application of electric fields to the culture, electrodes (5 cm; 0.22 mm^2^; stranded wire LiY; Conrad Electronic AG) were placed through the chamber-channels and adjusted to each end of the observation window at a distance of 3 cm. Electric fields up to 11 V/cm were applied using a low-voltage power supply (NTN 140-35; FuG Elektronik GmbH; 0-35V (±0.1V)) connected by crocodile clips (1m; MSL-100; Voltkraft® Conrad Electronic AG). Prior every experiment the chamber was washed with analytical grade Milli-Q water (Millipore Advantage system with Millipak Filter, Merck KGaA, Darmstadt, Germany) and the electrodes were used for no more than 10 electric field applications to minimize the oxidation effects on the electrode tips.

### Time-lapse microscopy

Time-lapse records were obtained by a stereo microscope (Stemi 508, Zeiss, Carl Zeiss Microscopy GmbH) equipped with a camera (Axiocam 503, Zeiss, Carl Zeiss Microscopy GmbH). The spatial magnification was set to 20.475× by using a 0.65× zoom, a 63× objective lens and a 0.5×wide angle camera adapter. Time-series were taken by using the ZEN software (ZEN lite - blue edition; Carl Zeiss Microscopy GmbH) applying an exposure time of 42 ms and spatial cropping of 1920×400 pixels, which led to a spatiotemporal resolution of 21 frames per seconds and 4.54 µm per pixel.

### Light microscopy

Single cell images were obtained by a ZEISS Axio Scope.A1 microscope equipped with a Zeiss EC Plan-Neofluar 63×/1,25 Oil M27 objective (Carl Zeiss Microscopy GmbH, Jena, Germany) equipped with blue light filter (Carl Zeiss Microscopy GmbH, Jena, Germany) using the differential interference contrast (DIC) mode. Images were recorded at 1.936 × 1.460 pixels with a ZEISS Axiocam 503 color Digital Camera and recorded using the Zeiss ZEN software (Vers. 2.3 blue edition; Carl Zeiss Microscopy GmbH, Jena, Germany). The movie was recorded with the Kinetix sCMOS Camera (Photometrics) with spatiotemporal resolution of 8 bit with 1000 frames per second and 0.10 µm per pixel.

### Scanning electron microscopy

Critical point dried samples of *C. hirtus* cells were imaged by scanning electron microscope (SEM). For this 50 ml of *C. hirtus* culture was concentrated by centrifugation (1121 rpm, 2 min, Sigma 6-10) in an oil test cup. The cells were fixed at room temperature for approx. 10 min by adding 100 µl 4 % osmium (OSO4)) solution to 1 ml of the cell sample. The cells were placed on a filter (25 mm, 0.4 µm, Nuclepore Track-Etch Membrane, Whatman) and dehydrated with 2,2-dimethoxipropane (Sigma-Aldrich). Subsequently, the 2,2-dimethoxipropane was replaced by liquid CO_2_ in 5 cycles and the cells were critically dried (CPD 020; Balzer Union). The filter was fixed onto aluminum specimen stubs covered with carbon Leit-tabs (12mm diameter, Plano GmbH, Germany) and sputtered with gold (31.8 mA, 70 s, 1.7 × 10-2 bar, MED 020 Coating System; BAL-TEC). Scanning electron microscopy images were obtained at 15 kV by a Zeiss EVO 15 scanning electron microscope (SEM) equipped with the Smart SEM software Ver. 6.04 SP3 (Zeiss, Oberkochen, Germany) using the secondary electron detector.

### Image Processing

All image series were analysed using the plugin TrackMate (v.5.02 (38)) in ImageJ (ImageJ 2.0 Fiji). Images were converted to 8 bit and the intensity inverted prior analysis. The DoG detector with the Simple LAP tracker was used to extract the spatiotemporal positions of the cells (see supplemental materials for details). The further statistical analysis of the extracted data was performed using customized routines in Matlab (R2020a; The MathWorks, Natick, MA).

### Statistical Analysis

The statistical analysis of the standard deviation of the velocities of individual cells 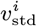 were log-transformed, because of the strongly skewed normal distributions, as

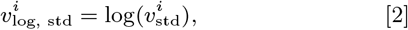

the mean *µ*_log_ and standard deviation *σ*_log_ was calculated, as

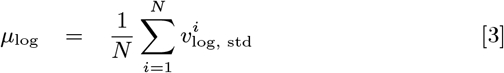

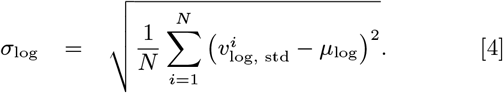

and back-transformed to obtain the mean and variance in the normal space as

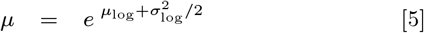

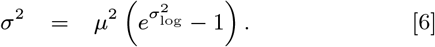

following the Finney estimator approach (39).

The direction probability *p* was fitted by least-square fits using normal cumulative distribution functions, as

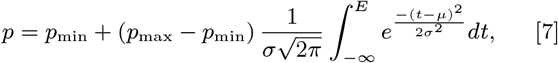

where *p*_min_ and *p*_max_ are the lower and upper bounds of *p, µ* and *σ* the mean and standard deviation of the underlying normal distribution, and the electric field *E* the depending parameter, which is parameterized by *t* in the interval (−∞, *E*].

The mean velocity variability ⟨*v*_std_⟩ was fitted by a piecewise logarithmic function as

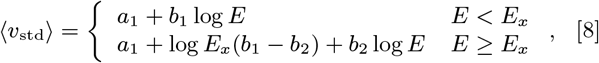

that fulfills the condition

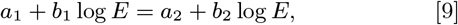

so that the function is continuous at the breakpoint *E*_*x*_ with the parameters *a*_*i*_ and *b*_*i*_.

## ACKNOWLEDGMENTS

We thank D. Walz for providing the 3D printed chamber. We also acknowledge S. Schmiech, A. Majoul and J. Reuter for their help with the cell cultures in the laboratory. We thank PD Dr. M. Schweikert and Prof. Dr. I. Weiss for the stimulating discussions and the critical reading of the manuscript. This research was supported by the German Research Foundation (DFG, AOBJ: 642944) and the European Regional Development Fund (EFRE, FEIH_778511).

