## Supplemental File for "Galvanotaxis of ciliates: Spatiotemporal dynamics of *Coleps hirtus* under electric fields"

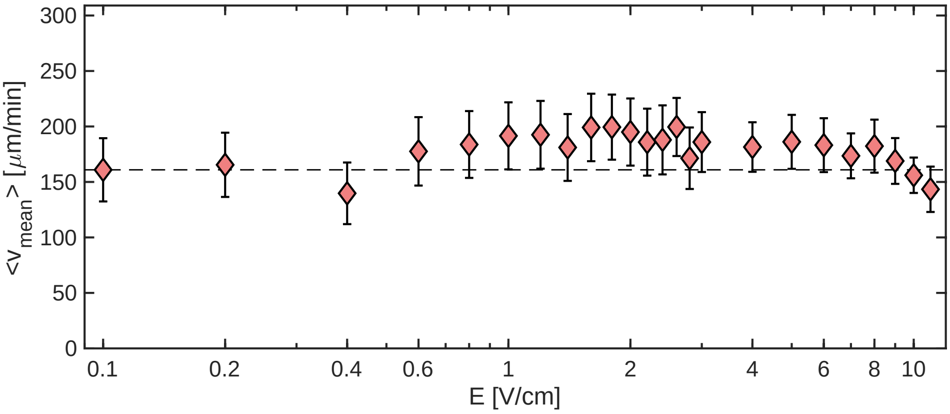


Figure S1: Mean velocity as a function of the electric field strength. The dashed line show the mean velocity of cells without the application of an electric field ($E = 0$ V/cm). Errorbars show the standard error.

### Cell medium preparation

Wright's cryptophytes (WC) medium was prepared by devolving $Na_2EDTA \times 2\,H_2O$, $NaHCO_3$, $NH_4Cl$, $CaCl_2 \times 2\,H_2O$, $K_2HPO_4 \times 3\,H_2O$, $MgSO_4 \times 7\,H_2O$, $NaNO_3$ and $H_3BO_3$ in 900 ml analytic-$H_2O$ (Table 1). Additionally 250 µL metal and 5 mL soil-trunk solution was added and the pH was adjusted to 7.8±0.2 using 0.5M NaOH and 0.5M HCl. The solution was centrifuged (10 min, 4000 rpm, RT), the volume of the medium was adjust to 1 L, autoclaved, and 500 µL vitamin solution added under steril conditions. The WC-medium was stored at 4°C until use.

Table 1: Wright's cryptophytes (WC) medium

| | **Stock solution** [g/L] | **Volume** [µL] |
|---|---|---|
| $Na_2EDTA \times 2\,H_2O$ | 37.2 | 50 |
| $NaHCO_3$ | 12.6 | 50 |
| $NH_4Cl$ | 26.8 | 100 |
| $CaCl_2 \times 2\,H_2O$ | 36.76 | 100 |
| $K_2HPO_4 \times 3\,H_2O$ | 8.71 | 250 |
| $MgSO_4 \times 7\,H_2O$ | 36.97 | 250 |
| $NaNO_3$ | 85.01 | 250 |
| $H_3BO_3$ | 6.0 | 250 |

The stock of the metal solution was prepared by dissolving 1.5 mL of each of the four stock solutions I - IV (Table 2) in 100 mL analytic-$H_2O$, adjusting the pH to 3.2 using 0.5M NaOH, adding 150 mL analytic-$H_2O$ and sterile filtration.

Table 2: Metal solution for WC-medium

| **Parts** | **Stock solution** | **Mass of chemicals** [mg] | **Volume of $H_2O$** [mL] |
|---|---|---|---|
| I | $ZnSO_4 \times 7\,H_2O$ | 175 | 50 |
| | $CuSO_4 \times 5\,H_2O$ | 30 | 50 |
| | $CoSO_4 \times 7\,H_2O$ | 60 | 50 |
| | $MnSO_4 \times H_2O$ | 61 | 50 |
| | → mix and filter sterile | | |
| II | $FeCl_3 \times 6\,H_2O$ | 300 | 6 |
| III | $Na_2MoO_4 \times 2\,H_2O$ | 3.3 | 100 |
| IV | $Na_2EDTA \times 2\,H_2O$ | 250 | 100 |

The stock of the soil-trunk solution was prepared by autoclaving 100 g unfertilised bedding soil (loamy soil Hohenheim, Germany) with 735 mL deionised water for 2 hours. The solution was allowed to settle and cool down overnight, and filtered through folded filters on the next day.

The vitamin solution was prepared by dissolving the following vitamins B1, B7 (Biotin), and B12 (see Table 3) in 100mL analytic-$H_2O$ and sterile filtration.

Table 3: Vitamin solution for WC-medium

| Vitamin | amount for stock | Mass of vitamin [mg] | Volume of $H_2O$ [mL] |
|---|---|---|---|
| B1 | 20 mg | | |
| B7 | 1.0 mL | 1 | 10 |
| B12 | 0.1 mL | 10 | 10 |

### Image analysis settings

For analysing the recorded time-series the plugin TrackMate (v.5.02 (38)) in ImageJ (ImageJ 2.0 Fiji) was used. First original tiff-stacks were loaded, the colormap (intensities) inverted, and the images to 8 bit converted. The following workflow was performed using the plugin:

1. [**Settings:**] Calibration & Crop settings (no changes)
2. [**Detector**] → DoG detector
3. [**DoG detector**]
   → Estimated blob diameter: 10.0 px
   → Threshold: 1.0
   → Use median filter: no
   → Do sub-pixel localization: no
4. [**Initial Thresholding**] → (no changes)
5. [**Select a view**] → hyperstack displayer
6. [**Set filters on spots**] (none)
7. [**Select a tracker**] → simple LAP tracker
8. [**Simple LAP tracker** ]
   → Linking max distance: 15.0 px
   → Gap-closing max distance: 15.0 px
   → Gap-closing max frame gap: 2
9. [**Set filters on track**] (none)
10. [**Display options**] → Analysis (save resulting files)

Further analysis of the saved output was performed by customized Matlab routines.
